# Common homozygosity for predicted loss-of-function variants reveals both redundant and advantageous effects of dispensable human genes

**DOI:** 10.1101/819615

**Authors:** A Rausell, Y Luo, M Lopez, Y Seeleuthner, F Rapaport, A Favier, PD Stenson, DN Cooper, E Patin, JL Casanova, L Quintana-Murci, L Abel

## Abstract

Humans homozygous or hemizygous for variants predicted to cause a loss of function of the corresponding protein do not necessarily present with overt clinical phenotypes. However, the set of effectively dispensable genes in the human genome has not yet been fully characterized. We report here 190 autosomal genes with 207 predicted loss-of-function variants, for which the frequency of homozygous individuals exceeds 1% in at least one human population from five major ancestry groups. No such genes were identified on the X and Y chromosomes. Manual curation revealed that 28 variants (15%) had been misannotated as loss-of-function, mainly due to linkage disequilibrium with different compensatory variants. Of the 179 remaining variants in 166 genes (0.82% of 20,232 genes), only 11 alleles in 11 genes had previously been confirmed experimentally to be loss-of-function. The set of 166 dispensable genes was enriched in olfactory receptor genes (41 genes), but depleted of genes expressed in a wide range of organs and in leukocytes. The 125 dispensable non-olfactory receptor genes displayed a relaxation of selective constraints both between species and within humans, consistent with greater redundancy. In total, 62 of these 125 genes were found to be dispensable in at least three human populations, suggesting possible evolution toward pseudogenes. Out of the 179 common loss-of-function variants, 72 could be tested for two neutrality selection statistics, and eight displayed robust signals of positive selection. These variants included the known *FUT2* mutant allele conferring resistance to intestinal viruses and an *APOL3* variant involved in resistance to parasitic infections. Finally, the 41 dispensable olfactory receptor genes also displayed a strong relaxation of selective constraints similar to that observed for the 341 non-dispensable olfactory receptor genes. Overall, the identification of 166 genes for which a sizeable proportion of humans are homozygous for predicted loss-of-function alleles reveals both redundancies and advantages of such deficiencies for human survival.

**Significance statement:** Human genes homozygous for seemingly loss of function (LoF) variants are increasingly reported in a sizeable proportion of individuals without overt clinical phenotypes. Here, we found 166 genes with 179 predicted LoF variants for which the frequency of homozygous individuals exceeds 1% in at least one of the populations present in databases ExAC and gnomAD. This set of putatively dispensable genes showed relaxation of selective constraints suggesting that a large number of these genes are undergoing pseudogenization. Eight of the common LoF variants displayed robust signals of positive selection including two variants located in genes involved in resistance to infectious diseases. The identification of dispensable genes will allow identifying functions that are, at least nowadays, redundant, or possibly advantageous, for human survival.

## Introduction

The human genome displays considerable DNA sequence diversity at the population level. One of its most intriguing features is the homozygosity or hemizygosity for variants of protein-coding genes predicted to be loss-of-function (LoF) found at various frequencies in different human populations (1–3). An unknown proportion of these reported variants are not actually LoF, instead being hypomorphic or isomorphic, because of a re-initiation of translation, readthrough, or a redundant tail, resulting in lower, normal, or even higher than normal levels of protein function. Indeed, a *bona fide* nonsense allele, predicted to be LoF, can actually be gain-of-function (hypermorphic), as illustrated by I*κ*B*α* mutations (4). Moreover, the LoF may apply selectively to one or a subset of isoforms of a given gene, but not others (e.g. if the exon carrying the premature stop is spliced out for a specific set of alternative transcripts) (5). Finally, there are at least 400 discernible cell types in the human body (6), and the mutant transcript may be expressed in only a limited number of tissues. Conversely, there are also mutations not predicted to be LoF, such as in-frame insertions-deletions (indel), missense variants, splice-region variants affecting the last nucleotides (nt) of exons and even synonymous or deep intronic mutations, that may actually be LoF but cannot be systematically identified as such *in silico*.

Many predicted LoF variants have nevertheless been confirmed experimentally, typically by demonstration of their association with a clinical phenotype. Of the 229,161 variants reported in HGMD (7), as many as 99,027 predicted LoF alleles in 5,186 genes have been found to be disease-causing in association and/or functional studies. For example, for the subset of 253 genes implicated in recessive forms of primary immunodeficiencies (8), 12,951 LoF variants are reported in HGMD. Conversely, a substantial proportion of genes harboring biallelic null variants have no discernible associated pathological phenotype, and several large-scale sequencing surveys in adults from the general population have reported human genes apparently tolerant to homozygous LoF variants (9–14). Four studies in large bottlenecked or consanguineous populations detected between 781 and 1,317 genes homozygous for mutations predicted to be LoF (10,11,13,14). These studies focused principally on low-frequency variants (minor allele frequency, MAF<1%-5%), and associations with some traits, benign or disease-related, were found for a few rare homozygous LoF variants. However, two studies provided a more comprehensive perspective of the allele frequency spectrum of LoF variants. A first systematic survey of LoF variants, mostly in the heterozygous state, was performed with whole-genome sequencing data from 185 individuals of the 1000 Genomes Project; it identified 253 genes with homozygous LoF variants (9). In a larger study of more than 60,000 whole exomes, the ExAC project identified 1,775 genes with at least one homozygous LoF variant, with a mean of 35 homozygous potential LoF variants per individual (12).

These studies clearly confirmed the presence of genes with homozygous LoF variants in apparently healthy humans, but no specific study of such variants present in the homozygous state in a sizeable proportion (>1%) of large populations has yet been performed. In principle, these variants may be neutral (indicating gene redundancy) (15), or may even confer a selective advantage (the so-called “less is more” hypothesis) (3, 16). Indeed, a few cases of common beneficial LoF variants have been documented, including some involved in host defense against life-threatening microbes (17, 18). Homozygosity for LoF variants of *DARC (*now *ACKR1), CCR5*, and *FUT2* confer resistance to *Plasmodium vivax* (*19, 20*), HIV (21–23), and norovirus (24, 25), respectively. We hypothesized that the study of common homozygous LoF variants might facilitate the identification of the set of dispensable protein-coding genes in humans and reveal underlying evolutionary trends. Unlike rare LoF variants, *bona fide* common homozygous LoF variants are predicted to be enriched in neutral alleles (1–3). They also provide indications concerning genes undergoing inactivation under positive selection (17, 18). The availability of large public databases, such as ExAC (12), and its extended version gnomAD, which includes data for more than 120,000 individuals (26), is making it possible to search for such variants with much greater power, across multiple populations.

## Results

### Definition of the set of dispensable protein-coding genes in humans

We used two large exome sequence databases: the Exome Aggregation Consortium database (ExAC, http://exac.broadinstitute.org/, (12)) and the Genome Aggregation Database (GnomAD, http://gnomad.broadinstitute.org/), which have collected 60,706 and 123,136 exome sequences, respectively. For this study, we focused on the 20232 protein-coding genes and we excluded the 13,921 pseudogenes (Methods; (27)). We defined protein-coding genes as dispensable if they carry variants that: (i) are computationally predicted to be loss-of-function (LoF) with a high degree of confidence, including early stop-gains, indel frameshifts, and essential splice site-disrupting variants (i.e. involving a change in the 2 nt region at the 5’ or 3’ end of an intron; **Methods**); and (ii) have a frequency of homozygous individuals (hemizygous for genes on the X chromosome) exceeding 1% in at least one of the five population groups considered in these public databases (i.e. Africans, including African Americans, East Asians, South Asians, Europeans, and Admixed Latino Americans). Common quality filters for calls and a minimum call rate of 80% were applied to each reference database (ExAC and GnomAD; **Methods**). Only LoF variants affecting the principal isoform (28) were retained (**Methods**). The application of these filters led to the detection of 208 (ExAC) and 228 (GnomAD) genes, 190 of which common to the two databases, and are referred to hereafter as the *set* of dispensable genes (**Table 1** and **Supplementary Table 1**). No genes on the X or Y chromosomes fulfilled these criteria. Relaxing the thresholds on the SNP and INDEL call quality filters (variant quality score recalibration (VQSR) score; **Methods**) or the variant call rate did not substantially increase the number of putatively dispensable genes (**Supplementary Figures 1** and **2**). The frequency of homozygous individuals at which a gene is defined as dispensable appeared to be the criterion with the largest impact on the number of dispensable genes identified, thereby justifying the use of the stringent threshold (>1% homozygotes in at least one specific population) described above (**Supplementary Figures 1** and **2**).

**Table 1.**
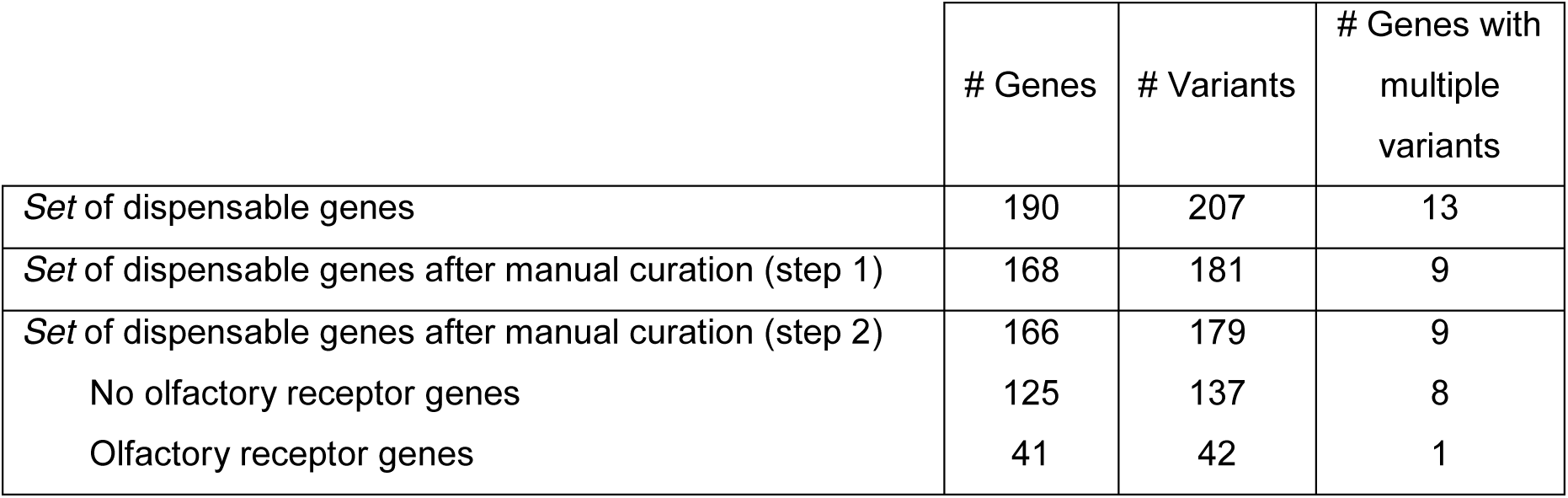
Number of dispensable genes identified depending on the adopted filtering criteria.

### Manual curation of the LoF variants

We then investigated the potential molecular consequences of the 207 putative LoF variants retained, collectively associated with the 190 genes, by manually curating the annotation of these LoF variants. An examination of the sequencing reads in GnomAD showed that 18 variants — 10 single nucleotide variants (SNVs) and eight indels initially annotated as stop-gains and frameshift variants, respectively — were components of haplotypes with consequences other than the initial LoF annotation (**Supplementary Table 1**). Thus, for each of the 10 SNVs, a haplotype encompassing a contiguous second variant led to the creation of a missense rather than a stop variant allele. For each of the eight frameshift variations, a haplotype with a nearby second indel (observed in the same sequencing reads) collectively led to an in-frame indel allele. In addition, six essential splice site-disrupting variants caused by indels resulted in no actual modification of the splice receptor or acceptor site motif. Overall, annotation issues occurred in about 13% of the initial set, indicating that there is a need for annotation methods to take the underlying haplotype inferred from sequencing reads into account. We also analyzed the common putative *HLA-A* LoF frameshift variant rs1136691 in more detail, as *HLA-A* null alleles are known to occur very rarely in large transplantation databases (http://hla.alleles.org/alleles/nulls.html). An analysis of the sequencing reads corresponding to this variant revealed an alternative haplotype of several variants, and a Blast analysis of this sequence yielded a perfect match with an alternative unmapped contig of chromosome 6 (chr6_GL000256v2_alt in GRCh38). This observation suggests that the alternative haplotype may have been wrongly mapped to the closest sequence in the reference genome, resulting in an artefactual frameshift in the *HLA-A* gene. This hypothesis is consistent with the known genomic complexity and high level of polymorphism of the *HLA* region, which can lead to incorrect mapping (29). Finally, we noted that one variant, rs36046723 in the *ZNF852* gene, had an allele frequency >0.9999, suggesting that the reference genome carries the derived, low-frequency variant at this position. After curation, we retained 181 predicted LoF variants of 168 genes (**Table 1**).

### Characteristics of the LoF variants

The set of 181 predicted LoF variants defining the set of dispensable genes included premature stop-gains (40%), frameshifts (47%), and splice-site variants (13%; **Figure 1**). The variants could be classified further into those with a high or low predicted probability of being LoF (**Methods**): 27% presented features consistent with a low probability of LoF, whereas the remaining **73**% were predicted to have more severely damaging consequences and were therefore considered to have a high probability of LoF (**Figure 1** and **Supplementary Figure 3**). Despite possible differences in their impact, low and high probability LoF variants had similarly distributed global allele frequencies (**Supplementary Figure 4**; two-tailed Wilcoxon test *p*-value = 0.3115, based on ExAC allele frequencies; and 0.3341 based on GnomAD allele frequencies). Only 30 of the LoF variants finally retained were reported in at least one PubMed publication in the dbSNP database (30) (**Supplementary Table 1**). Manual inspection of these studies revealed that only 11 LoF variants had been experimentally demonstrated to abolish gene function (**Supplementary Table 1**). Focusing on the overlap between the 181 predicted LoF variants and the GWAS hits, we found that only two were reported in the GWAS catalog (31) as being significantly associated with a phenotypic trait: rs41272114 (*LPA*), associated with plasma plasminogen levels, lipoprotein A levels and coronary artery disease, and rs601338 (*FUT2*), associated with the levels of certain blood proteins, such as fibroblast growth factor 19. Overall, this analysis shows that most of the common LoF considered here present features consistent with severely damaging variants, although experimental characterization is largely lacking.

**Figure 1.**
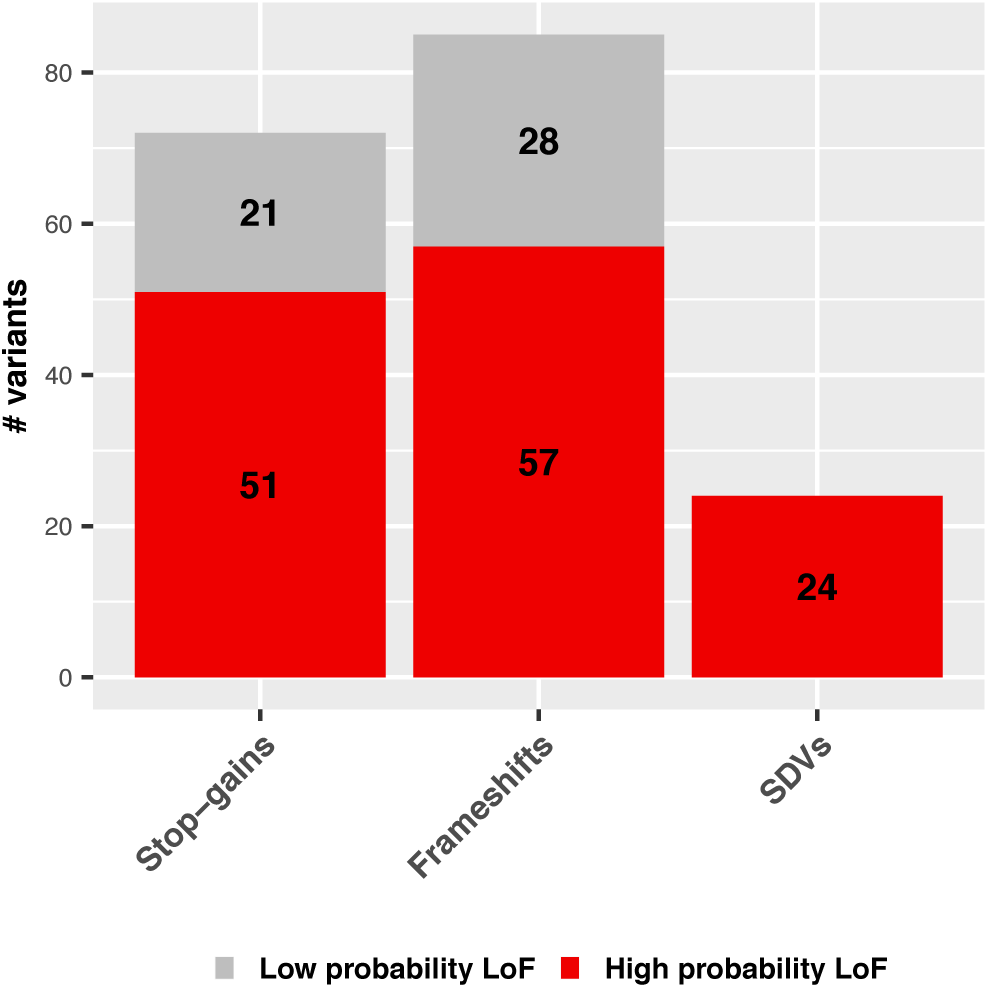
Functional consequences of LoF variants defining the set of dispensable protein-coding genes. Barplots show the distribution of LoF variants defining the set of dispensable genes according to their molecular consequences (stop-gains, frameshifts and splice-disrupting variants, SDVs) and the predicted severity of their functional impact: probably low probability (light grey) and high probably LoF (dark red; **Methods**).

### Overlap of dispensable genes with disease-causing and essential genes

We then focused on the features characterizing the list of 168 putatively dispensable genes, and searched for those previously shown to be (i) associated with Mendelian diseases (*n*=3,314, OMIM (32)), or (ii) essential in human cell lines (*n*=2,010) or knockout mice (*n*=3,326) (33). We focused on LoF variants predicted to be severely damaging, as described above (**Figure 1**). We found that three LoF variants from our list affected OMIM genes (*CLDN16, TMEM216, GUF1;* **Table 2**), while none affected essential genes (Fisher’s exact p-value = 7.727e-06 and 6.249e-10 for human cell lines and knockout mice essential genes, respectively). It has been suggested that the common LoF variant rs760754693 of *CLDN16* — a gene associated with a renal disorder known as familial hypomagnesemia with hypercalciuria and nephrocalcinosis — affects the 5’-untranslated region of the gene rather than its coding sequence (34). A second methionine residue downstream of the affected position in *CLDN16* could potentially serve as the actual translation start site. The *TMEM216* variant rs10897158 is annotated as benign in ClinVar (35). *TMEM216* is required for the assembly and function of cilia, and pathogenic mutations of this gene cause Joubert, Meckel and related syndromes (36). The canonical transcript of *TMEM216* encodes a 145-amino acid protein. The rs10897158 splice variant (global frequency of homozygous individuals >70%) results in the synthesis of a longer protein (148 amino acids) corresponding to the most prevalent isoform (36), which could probably be considered the reference protein in humans. There is currently little evidence to support the third variant, rs141526764 (*GUF1*), having any pathogenic consequences. The association of *GUF1* with Mendelian disease (early infantile epileptic encephalopathy) is reported in OMIM as provisional and based solely on the finding of a homozygous missense variant (A609S) in three siblings born to consanguineous parents (37). For all subsequent analyses, we filtered out the two variants considered highly likely not to be LoF (rs760754693, rs10897158), resulting in a final list of 179 predicted LoF variants corresponding to 166 genes (**Table 1**). The absence of common LoF variants predicted with a high degree of confidence in disease genes or essential genes is consistent with these genes being dispensable.

**Table 2.** Common high-probability LoF variants in OMIM genes.

| Gene | rsID | Consequence | Global allele Frequency (GnomAD) | % of homozygous individuals per population (GnomAD) |
| --- | --- | --- | --- | --- |
| *CLDN16* | rs760754693 | Frameshift | 0.194 | AFR (0.68%)<br>AMR (1.50%)<br>EAS (0.05%)<br>EUR (6.07%)<br>SAS (3.18%) |
| *GUF1* | rs141526764 | Splice Donor Variant | 0.015 | AFR (0.00%)<br>AMR (1.35%)<br>EAS (0.00%)<br>EUR (0.00%)<br>SAS (0.00%) |
| *TMEM216* | rs10897158 | Splice Acceptor Variant | 0.830 | AFR (12.83%)<br>AMR (74.35%)<br>EAS (92.64%)<br>EUR (72.84%)<br>SAS (70.05%) |

### Features of the set of putative dispensable protein-coding genes

We characterized the set of 166 putatively dispensable protein-coding genes further by performing a Gene Ontology (GO) enrichment analysis (**Methods**). We found a significant overrepresentation of genes involved in G-protein coupled receptor activity and related GO terms (Fisher’s *p*-value= 5.50e-25; **Supplementary Table 2**). Such enrichment was driven by the presence of 41 olfactory receptor (OR) genes, accounting for 25% of the total set of dispensable genes, consistent with previous analyses (9). After the removal of ORs, a few functional categories were identified as displaying significant enrichment (see **Supplementary Table 2** for the full list of enrichments). Proteins associated with keratin intermediate filaments or ligand-gated ion channel activity were the only top-scoring functional terms displaying enrichment among dispensable genes, but their uncorrected *p*-values were somewhat high (>1e-05). Thus, a large number of protein-coding genes may be dispensable, but this has no apparent impact at the level of pathways or functional categories. Based on previous findings and the known specific features of OR genes (38, 39), we decided to consider the 41 OR and 125 non-OR dispensable genes separately in subsequent analyses (**Table 1**). In comparisons with a reference set of 382 OR, and 19850 non-OR protein-coding genes (Methods), the coding lengths of the 41 OR (median = 945 nt) and the 125 non-OR (median = 1153.5 nt) dispensable genes were not significantly different from that of the corresponding non-dispensable OR (median = 945 nt; Wilcoxon test *p*-value= 0.6848) and non-OR (median = 1275 nt; Wilcoxon test *p-*value=0.1263) genes. The genomic distribution of dispensable OR genes displayed some clustering on some chromosomes, but did not differ significantly from that of the reference OR genes (**Supplementary Figure 5A**). The distribution of dispensable non-OR genes across autosomal chromosomes was also similar to that for the reference set, except that no dispensable genes were present on the X and Y chromosomes (**Supplementary Figure 5B**). This finding suggests that common LoF variants on sexual chromosomes are more efficiently purged from the population than autosomal variants, possibly because they have pathogenic effects in hemizygous males.

### Organ expression patterns of the dispensable protein-coding genes

We then investigated the expression patterns of the 166 putatively dispensable genes across organs and leukocyte types. For organs, we used RNA-seq expression data from the Illumina Body Map project (IBM) and the GTEX project, and for leukocytes, we used data from the BluePrint project (**Methods; Supplementary Table 3**). Most of the dispensable OR (30 of 34, 88%) were not found to be expressed in any of the datasets considered, consistent with the general expression pattern for all OR genes (328 of 355 OR genes not expressed, 92%; **Figure 2**). We found that 996 of a reference set of 17948 non-OR protein-coding genes were not expressed in any of the databases considered (referred to hereafter as non-detected genes). Interestingly, the non-detected genes displayed a significant enrichment in dispensable genes relative to the reference set: odds ratio (OR): 3.34, 95% confidence interval (CI) 1.92-5.53, Fisher’s exact test *p*-value 2.29e-05 (**Figure 2**). A similar pattern was observed for organ-specific genes, which were defined as genes expressed in <20% of the organs evaluated in the corresponding dataset: OR of 2.09 (CI 1.38-3.11) for IBM, *p*-value=3.67e-04, and OR of 3.56 (CI 2.41-5.22) for GTEX, *p*-value=1.53e-10. We further characterized the distribution of non-OR dispensable genes among the organ-specific genes for the various organs evaluated. Consistent with previous observations (11), the brain appeared to be the only organ in which the proportion of dispensable genes was significantly lower than that observed for organ-specific genes in both the IBM and the GTEX datasets: OR of 0.23 (CI 0.04-0.72), *p*-value=5.47e-03, and OR of 0.08 (CI 0.002-0.45), *p*-value=2.69e-04, respectively (**Supplementary Table 4**). Organ-pervasive genes were also found to be significantly depleted of dispensable genes, reflecting a lower degree of redundancy: OR of 0.09 (CI 0.04-0.18), *p*-value=6.47e-20 for IBM, and OR of 0.16 (CI 0.09-0.26), *p*-value=8.04e-19 for GTEX. The number of organs in which a gene is expressed was consistently, and negatively associated with the proportion of dispensable genes (linear regression *p*-values after adjustment for coding sequence length < 2e-16 for both IBM and GTEX). Overall, genes widely expressed or specifically expressed in the brain are less dispensable than those with a more restricted pattern of expression.

**Figure 2.**
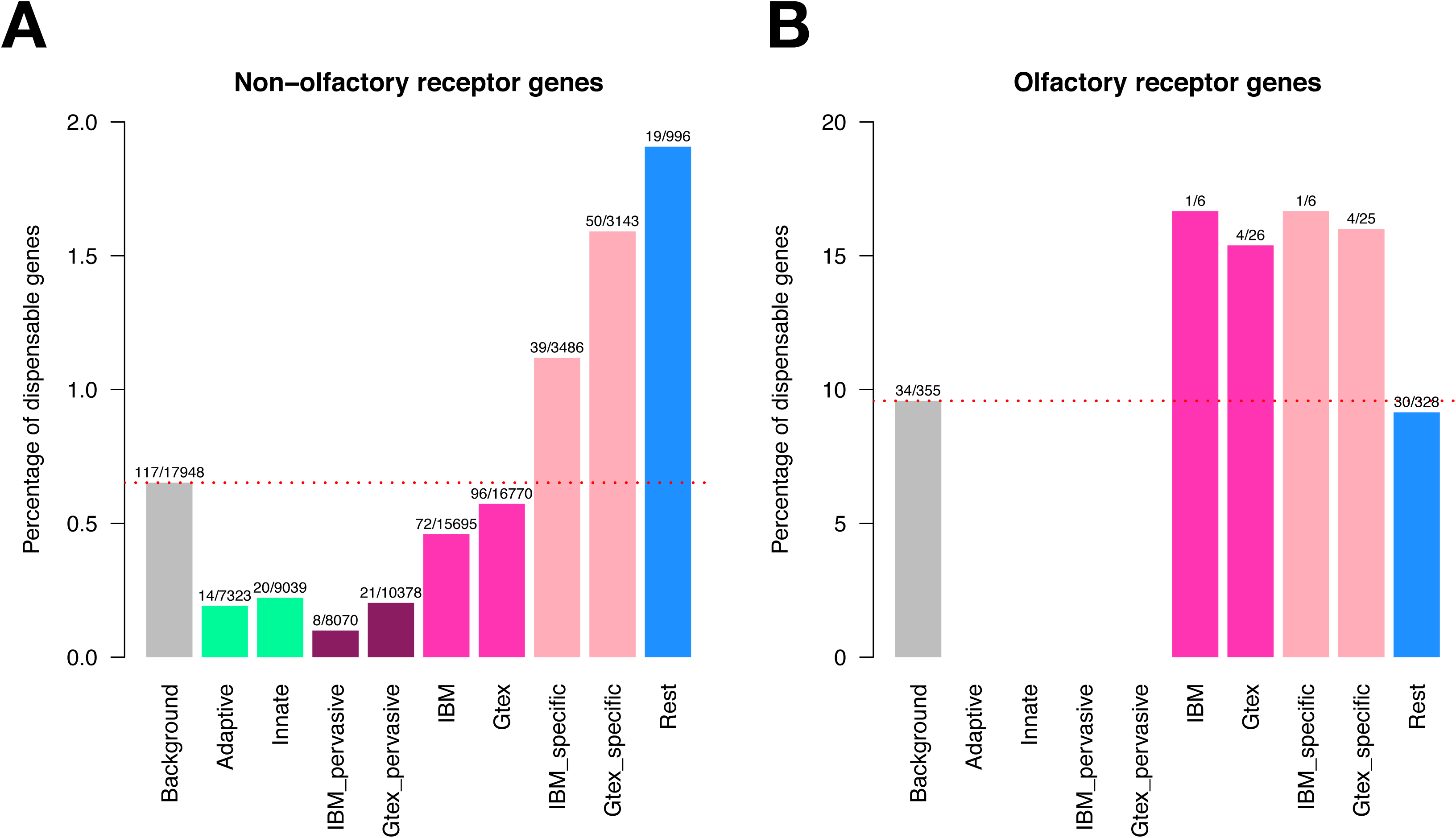
Distribution of the set of dispensable genes in tissue-expressed genes and adaptive and innate leukocytes expressed genes, defined from gene expression datasets. Relative enrichment in dispensable genes among different gene sets defined from expression datasets based on RNA-seq expression data from the Illumina Body Map project (IBM), the GTEX project and the BluePrint project (Adaptive and Innate leukocytes). Tissue-expressed genes are further classified in two subcategories: tissue-specific and tissue-pervasive genes (see text). The ratio of dispensable genes versus the total size of each category is indicated. Results are presented separately for 17948 non-OR genes (**A**) and 355 OR genes (**B**) for which expression data could be retrieved out of an initial list of 20232 protein-coding genes (**Methods**). P-values of the two-sided Fisher’s tests comparing the fraction of dispensable genes among the gene subsets against the reference background are reported in the text.

### Expression patterns for dispensable genes in leukocytes

We then investigated the expression patterns of the 166 putatively dispensable genes in leukocytes. Human immune genes were classified on the basis of the RNA-seq data generated by the BluePrint project (26). We identified 7,323 adaptive leukocyte-expressed genes on the basis of their expression in B or T cells in at least 20% of the samples considered, and 9,039 innate leukocyte-expressed genes defined on the basis of their expression in macrophages, monocytes, neutrophils, or dendritic cells (DC) in at least 20% of the samples considered (**Supplementary Table 3**). We are aware that the main function of DCs is to present antigens to T cells, making their classification as “innate” both arbitrary and questionable. These leukocyte-expressed genes included no OR genes, and all the results therefore correspond to non-OR genes. A significant depletion of dispensable non-OR genes relative to the reference set was observed among the genes expressed in adaptive and innate leukocytes: OR of 0.20 (95% CI 0.10-0.34), *p*-value=8.80e-12, and OR of 0.20 (CI 0.12-0.33), *p*-value=9.98e-14, respectively; **Figure 2**). In total, 24 dispensable genes were identified as expressed in adaptive leukocytes (*n*=4 genes), innate leukocytes (*n*=10) or both (*n*=10). Detailed information about the common homozygous LoF variants of these genes is presented in **Supplementary Table 5.** Sixteen of these 24 genes had variants predicted to have highly damaging consequences, including a well-known stop-gain variant of *TLR5* (40). This truncating *TLR5* variant, which abolishes cellular responses to flagellin, appears to evolve under neutrality (41). These genes also included *APOL3*, which is known to be involved in the response to infection with African trypanosomes (42). Thus, genes widely expressed in leukocytes are generally less dispensable than the reference set. However, specific immune-related genes may become LoF-tolerant due to functional redundancy (e.g. TLR5) or positive selection, by increasing protective immunity, for example. This aspect is considered in a subsequent section.

### Population distribution of the dispensable genes

We analyzed the distribution of the dispensable genes across the five specific populations considered: Africans (including African-Americans), East Asians, South Asians, Europeans (including Finnish), and Americans of Latino descent (**Figure 3**). Of the 125 non-OR genes, we found that 33 were dispensable in all populations, 16 in four populations, and 13 in three populations aggregated homozygous LoF frequency >1% in each population). Conversely, 48 genes were population-specific, with Africans providing the largest fraction in both absolute (26 genes) and relative terms, as a reflection of their greater genetic diversity (43). The remaining 15 genes were found in two populations. Almost half the 41 dispensable OR were common to all five populations (*n*=19), 10 were present in four or three populations, two were present in two populations, and 10 were population-specific (including nine in Africans). As expected, the number of populations in which a gene was found to be dispensable correlated with the maximum frequency of homozygous individuals in the populations concerned (**Supplementary Figure 6**). Overall, 49% of the non-OR and 71% of the OR dispensable genes (>90% of the dispensable OR genes in non-African populations) were common to at least three human populations, suggesting a general process of tolerance to gene loss independent of the genomic background of the population.

**Figure 3.**
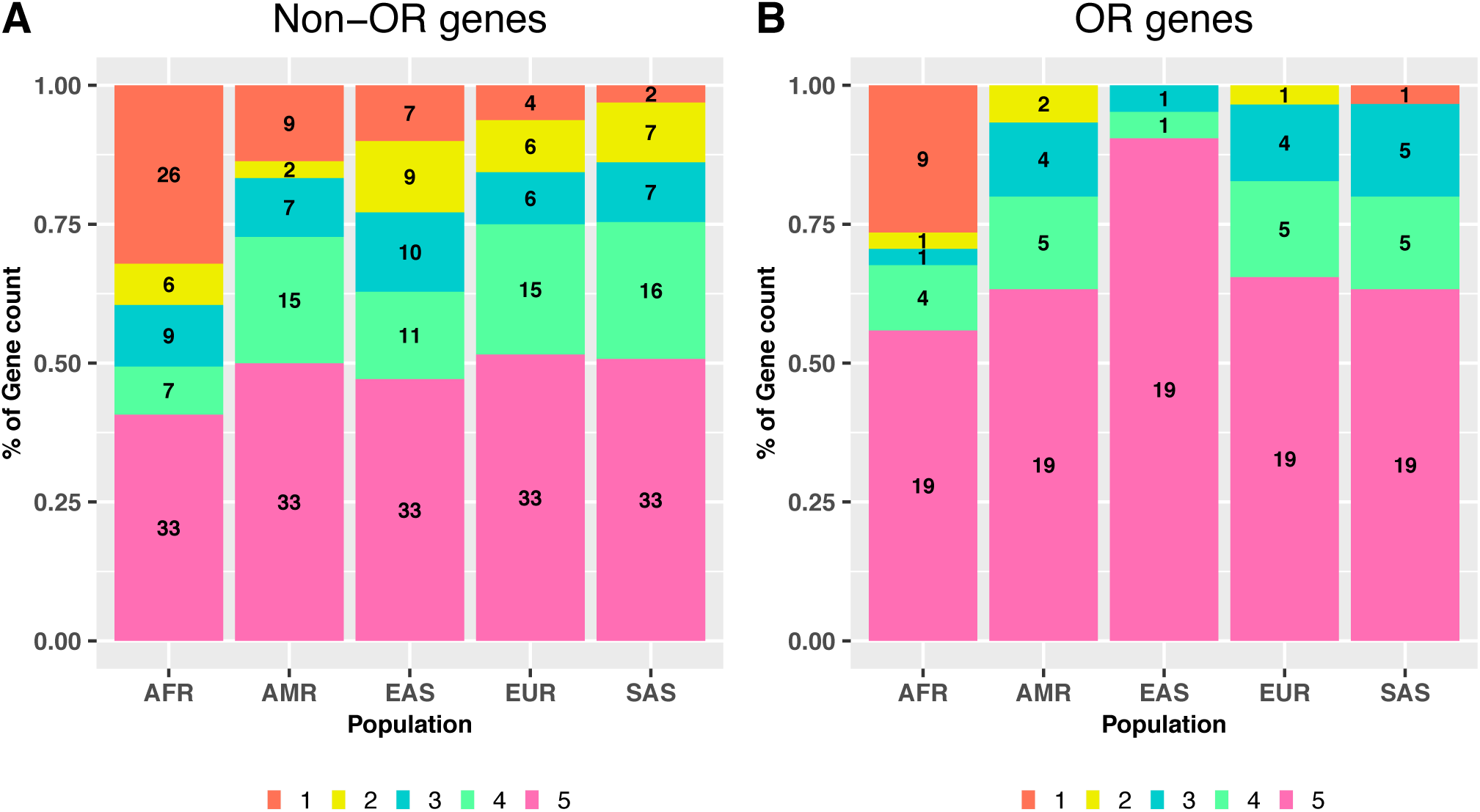
Distribution of dispensable genes across the 5 human populations considered. The number of dispensable non-OR genes (**A**) and dispensable OR genes (**B**) in each population (>1% aggregated homozygous LoF frequency in a given population) is represented across 5 categories indicating whether the gene is dispensable in all 5 populations considered, four, three or two of them, or is a population-specific dispensable gene. The five populations considered are: Africans (including African American, AFR), Americans (AMR), East Asians (EAS), Europeans (Finnish and Non-Finnish; EUR) and South Asians (SAS). The homozygous LoF variant frequencies were taken from GnomAD dataset for the purpose of this analysis.

### Negative selection of dispensable genes

We then investigated the behaviour of dispensable genes in terms of the functional scores associated with gene essentiality and selective constraints (**Figure 4**). We first evaluated a metric assessing the mutational damage amassed by a gene in the general population (GDI (44)) and three gene-level scores assessing recent and ongoing negative selection in humans (RVIS (45), pLI and pRec (46); **Methods**). Consistent with expectations, dispensable non-OR genes had much higher GDI (two-sided Wilcoxon test *p*-value=6.52e-35), higher RVIS (*p*-value=1.30e-37), and lower pLI (*p*-value=1.548597e-22) values than non-dispensable non-OR genes (**Figure 4**), whereas no significant differences were found for pRec values (*p*-value=0.29). In analyses focusing on OR genes, GDI and RVIS values were also significantly higher for dispensable genes than for non-dispensable genes (*p*-values= 3.08e-06 and 1.96e-02, respectively), whereas no differences were found for the pLI and pRec distributions (*p*-value>0.05). We then evaluated the strength of negative selection at a deeper evolutionary scale, using the estimated proportion *f* of non-synonymous mutations that are non-deleterious and have not, therefore, been purged from the population. Following an approach described elsewhere (47), *f* values were obtained with SnIPRE, by comparing polymorphism within humans and divergence between humans and chimpanzee at synonymous and non-synonymous sites (48). The *f* values obtained did not differ significantly (*p*-value >0.05) between dispensable and non-dispensable OR genes. However, dispensable non-OR genes had a distribution of *f* values reaching higher values than that of non-dispensable OR genes (*p*-value= 1.51e-29), indicating that dispensable OR genes were more tolerant of non-synonymous variants than non-dispensable OR genes. Dispensable genes also had more human paralogs than other protein-coding genes (*p*-value= 8.76e-06), suggesting a higher degree of redundancy. For OR genes, the number of paralogs did not differ between dispensable and non-dispensable genes, further confirming that the negative selection parameters of dispensable and non-dispensable genes OR genes were similar. Overall, these results reveal a relaxation of the selective constraints acting at dispensable non-OR gene loci relative to non-dispensable genes, providing further evidence for evolutionary dispensability.

**Figure 4.**
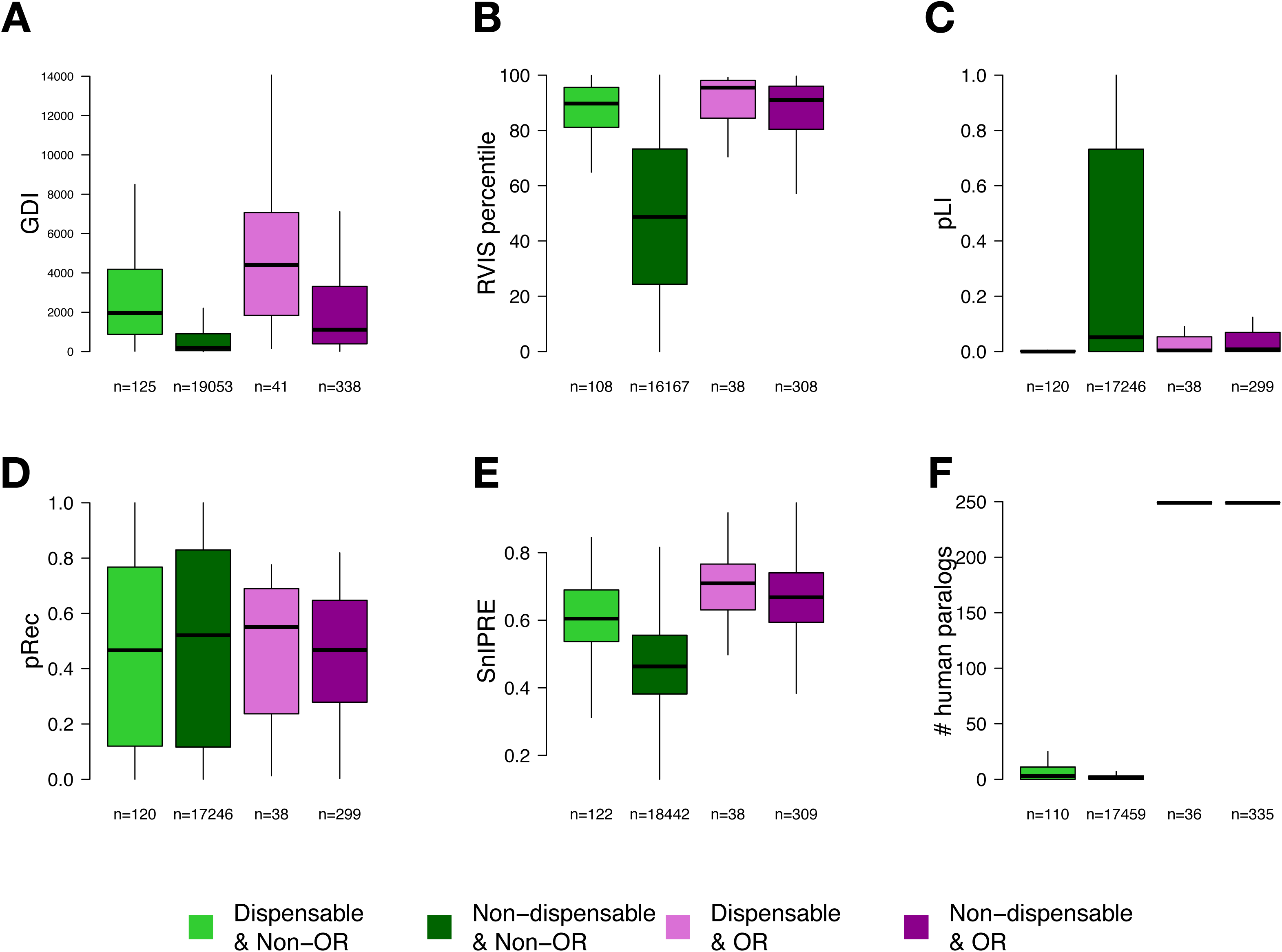
Distribution of functional scores related to gene essentiality/redundancy in the *stringent* set of dispensable genes. Score distributions presented include: **A**. Gene Damage Index (GDI). **B**. Residual Variation Intolerance Scores (RVIS). **C.** Probability of being intolerant to both heterozygous and homozygous LoF variants (pLI). **D.** Probability of being intolerant to homozygous LoF variants (pRec). **E**. SnIPRE values. **E.** Number of paralogs (**Methods**). Panels display the distribution of scores across dispensable non-OR genes (light green), non-dispensable non-OR genes (dark green), dispensable OR genes (light purple) and non-dispensable OR genes (dark purple). Two-sided Wilcoxon test p-values comparing the distribution of dispensable non-OR genes against non-dispensable non-OR genes are: (A) p-value=6.52e-35; (B) p-value=1.30e-37;(C) p-value=1.55e-22; (D) p-value=2.94e-01; (E) p-value=1.51e-29; (F) p-value=8.76e-06. Two-sided Wilcoxon test p-values comparing the distribution of dispensable OR genes against non-dispensable OR genes are: (A) p-value=3.04e-06; (B) p-value=1.96e-02; (C) p-value=4.64e-01; (D) p-value=7.98e-01; (E) p-value=6.81e-02; (F) p-value=1.23e-01.

### Positive selection of common LoF variants

We investigated the possibility that the higher frequency of some LoF mutations was due to a selective advantage conferred by gene loss (i.e., the “less-is-more” hypothesis), by searching for population-specific signatures of positive selection acting on these variants (17, 18). We considered two neutrality statistics: *F*_ST_, which measures between-population differences in allele frequencies at a given locus (49), and integrated haplotype score (iHS) (50), which compares the extent of haplotype homozygosity around the ancestral and derived alleles in a given population; both statistics could be computed for 72 variants fulfilling the quality control criteria (**Methods**). Considering a cutoff point of the 95^th^ percentile for each statistic (**Figure 5**), we detected 39 common LoF alleles in putatively dispensable genes displaying signals suggestive of positive selection, as attested by their low iHS (*n*=7), high *F*_ST_ values (*n*=24), or both (*n*=8; **Supplementary Table 6**). Seven of these variants in OR genes had only high *F*_ST_ values, suggestive of genetic drift related to the ongoing pseudogenization of ORs (39). We also noted that the LoF mutation (rs2039381) of *IFNE* displayed significant levels of population differentiation (e.g. *F*_ST_=0.25 for GIH vs. CEU, *P*_emp_ = 0.002). This result is intriguing as *IFNE* encodes IFNε, which plays an important role in protective immunity against microbes in the female reproductive tract in mice (51, 52). This nonsense variant (Q71X) is predicted to decrease the length of the encoded protein by two thirds, but this has not been validated experimentally. The proportion of homozygotes is highest in East Asia (3.5%) and South Asia (2%), is much lower in Europe (0.02%), and does not differ between males and females. The iHS scores were not significant for this variant (**Supplementary Figure 7**), and neither were other selection scores, such as Tajima’s *D*, Fu and Li’s *D*\* and *F* *, and Fay and Wu’s *H* t, previously obtained in a large evolutionary genetic study of human interferons (53). In light of these observations, the most likely explanation for the high *F*_ST_ observed at this locus is genetic drift.

**Figure 5.**
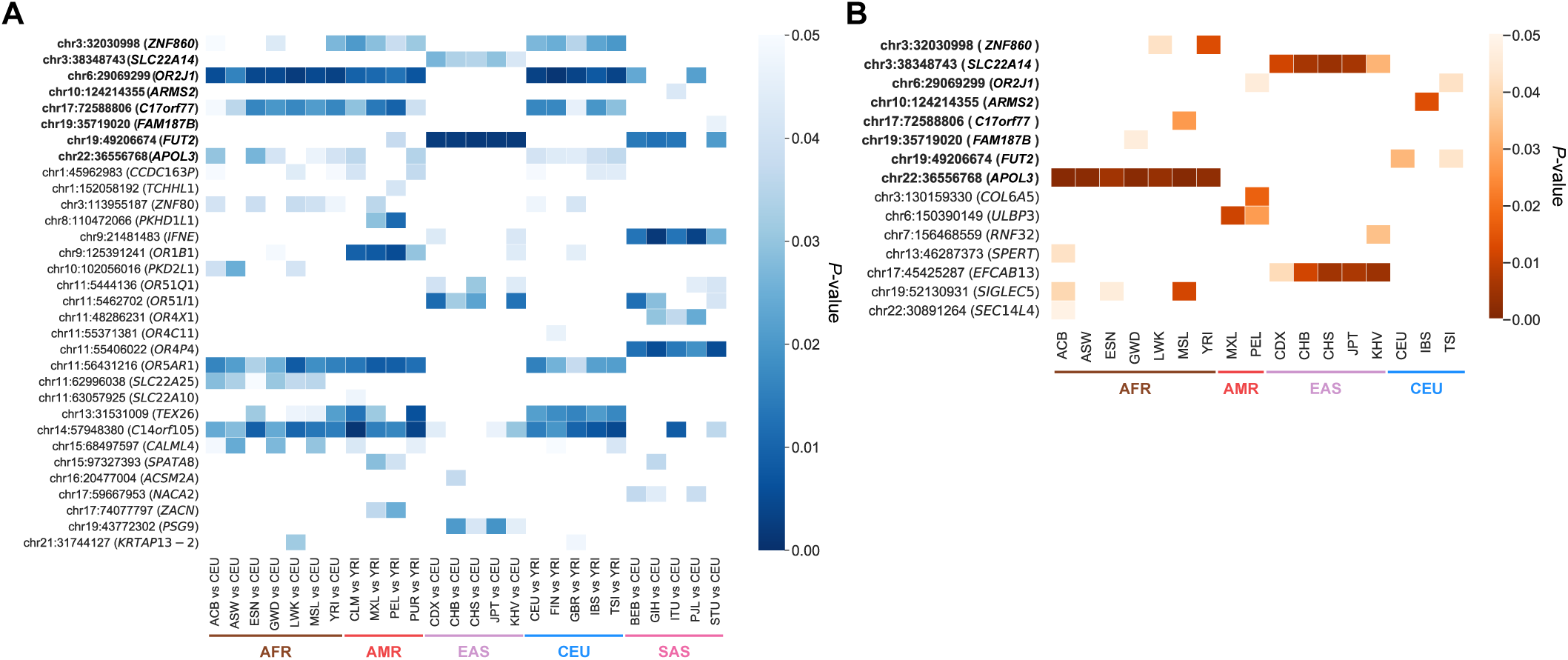
Evidence for positive selection on common LoF alleles. **A.** Empirical p-values for 32 LoF mutations presenting *F*_ST_ scores measuring allele frequency differentiation, in the 95^th^ percentile of highest values genome-wide, in at least one population. **B.** Empirical p-values for 15 LoF mutations presenting integrated haplotype scores (iHS) in the 95^th^ percentile of lowest values genome-wide (i.e. selection on the LoF allele), in at least one population. *F*_ST_ and iHS values are reported for each of the 26 populations of the 1000 Genomes Project, grouped in 5 super populations: AFR (brown), AMR (orange), EAS (purple), EUR (blue) and SAS (pink). Eight common LoF variants with both high *F*_ST_ and low iHS are listed first in bold. The colour gradients indicate the significance of the p-values and only polymorphic sites involving common biallelic LoF SNPs from the stringent set predicted to have severe damaging functional consequences were considered.

### LoF mutations of *FUT2* and *APOL3* are under positive selection

Eight common LoF variants provided more robust evidence for positive selection, as they had both a high *F*_ST_ and a low iHS (**Supplementary Table 6**, **Figure 5**). For five of these variants, there was no obvious relationship between the gene concerned and a possible selective advantage. One variant with a high *F*_ST_ (*F*_ST_ = 0.25 for ITU vs. CHB, *P*_emp_= 0.012) and a low iHS (iHS −2.33, *P*_emp_ = 0.004 in CHS) was located in *SLC22A14,* which has been shown to be involved in sperm motility and male infertility in mice (54). Finally, the two remaining variants were in the *FUT2* and *APOL3* genes, which are known to be involved in defense against infections. Consistent with previous observations, we observed high *F*_ST_ values (*F*_ST_ = 0.54 for CEU vs. CHS, *P*_emp_= 0.002) and extended haplotype homozygosity iHS =-1.7, *P*_emp_ = 0.03 in CEU) around the LoF mutation (rs601338) in *FUT2* (**Supplementary Figure 7**). This gene is involved in antigen production in the intestinal mucosa, and null variants are known to lead to the non-secretor phenotype conferring protection against common enteric viruses, such as norovirus (25, 55), and rotavirus (56, 57). We also identified a novel hit at the *APOL3* LoF variant rs11089781. This nonsense variant (Q58X) was detected only in African populations (15-33% frequency), in which it had a low iHS (iHS=-2.75 in MSL, *P*_emp_ = 0.001), indicating extended haplotype homozygosity around the LoF mutation in these populations (**Supplementary Figure 7**). *APOL3* is located within a cluster of *APOL* genes including *APOL1*. These two members of the six-member APOL cluster, APOL1 and APOL3, are known to be involved in defense against African trypanosomes (58, 42). These analyses indicate that, although positive selection remains rare in humans, it may have increased the frequency of LoF variants when gene loss represents a selective advantage.

## Discussion

We detected 166 putatively dispensable human protein-coding genes. These genes included 120 that overlapped either with the total list of 2,641 genes apparently tolerant to homozygous rare LoF reported from bottlenecked or consanguineous populations (10,11,13,14) or with the list of 253 genes initially identified from individuals of the 1000 Genomes Project (9) (**Supplementary Figure 8**), or both. Most of the 46 genes specific to our study had a maximum homozygote frequency below 0.05 or had a higher frequency only in one or two of the five studied populations, as for *FUT2* and *IFNE*. Our set of dispensable genes was strongly enriched in OR genes, as previously reported (9, 59). However, dispensable OR genes had no particular features distinguishing them from non-dispensable OR genes other than slightly higher GDI and RVIS values. This finding is consistent with the notion that ORs are increasingly becoming dispensable in the human lineage (60, 39). Conversely, dispensable non-OR genes displayed a strong relaxation of selective constraints relative to non-OR non-dispensable genes at both the inter-species and intra-species levels. In addition, the set of dispensable non-OR genes was depleted of genes widely expressed in the panel of organs evaluated, brain-specific genes and genes expressed in leukocytes. This suggests that the redundancy observed for some microbial sensors and effectors (61, 18) does not necessarily translate into higher rates of gene dispensability.

The set of dispensable genes identified here probably largely corresponds to genes undergoing pseudogenization (62, 63) due to present-day superfluous molecular function at the cell, organ or organism levels, or a redundant function in the genome that may be recovered (e.g. by paralogous genes or alternative pathways). This is the case for OR genes, which are strongly enriched in dispensable genes, and generally present a strong relaxation of selective constraints and signs of ongoing pseudogenization (39). Another example is provided by *TLR5,* encoding a cell-surface receptor for bacterial flagellin, which harbors a dominant negative stop mutation at high population frequencies (41,40,64). This finding is consistent with the notion that a substantial proportion of modern-day humans can survive with complete TLR5 deficiency (40). These observations also suggest that additional mechanisms of flagellin recognition, such as those involving the NAIP-NLRC4 inflammasome (65, 66), may provide sufficient protection in the absence of TLR5. Finally, 45 of our dispensable genes belong to the set of 2,278 Ensembl/GENCODE coding genes recently reported to display features atypical of protein-coding genes (67). Our study may therefore provide additional candidates for inclusion in the list of potential non protein-coding genes.

High population frequencies of LoF variants may also reflect recent and ongoing processes of positive selection favoring gene loss (i.e. the “less-is-more” paradigm) (17). Two of the eight variants with the most robust signals of positive selection were located in genes involved in resistance to infectious diseases. The *FUT2* gene is a well-known example of a gene for which loss is an advantage, as it confers Mendelian resistance to common enteric viruses and has a profile consistent with positive selection. An interesting new finding from this study is provided by the *APOL3* LoF variant (rs11089781), which we found to display signals of recent, positive selection in Africans. APOL3 and APOL1 are known to be involved in the response to African trypanosomes (42). Two variants encoding APOL1 proteins with enhanced trypanolytic activity are present only in African populations, in which they harbor signatures of positive selection despite increasing the risk of kidney disease (68). Interestingly, the *APOL3* LoF variant was also recently associated with nephropathy independently of the effect of the two kidney disease-risk *APOL1* variants, which are not in strong linkage disequilibrium with rs11089781 (69). A physical interaction occurs between APOL1 and APOL3 (69), and APOL1 may protect against pathogens more effectively when not bound to APOL3. Similar mechanisms may, therefore, be involved in the positive selection of the *APOL1* kidney disease-risk alleles and the *APOL3* LoF variant in African populations. Additional common LoF homozygotes could probably be further identified in other unstudied populations, or involving variants not currently predicted to be LoF *in silico*. Improvements in the high-confidence identification of dispensable genes will make it possible to identify functions and mechanisms that are, at least nowadays, redundant, or possibly advantageous, for human survival.

## Acknowledgments

We thank members of the Laboratory of Clinical Bioinformatics and both branches of the Laboratory of Human Genetics of Infectious Diseases, Yuval Itan, Sophie Saunier and Corinne Antignac for helpful discussions and support. The Laboratory of Clinical Bioinformatics was supported by grants from the French National Research Agency (Agence Nationale de la Recherche, ANR) “Investissements d’Avenir” program, (grant # ANR-10-IAHU-01) and the Christian Dior Couture, Dior. The Laboratory of Human Genetics of Infectious Diseases was supported in part by grants from ANR under the “Investissement d’avenir” program (grant # ANR-10-IAHU-01) and the “Laboratoires d’Excellence” Integrative Biology of Emerging Infectious Diseases (IBEID) (grant # ANR-10-LABX-62-IBEID), the French Foundation for Medical Research (FRM) (EQU201903007798), the National Center for Advancing Translational Sciences of the National Institutes of Health (grant # UL1TR001866), the St. Giles Foundation, and the Rockefeller University. The laboratory of Human Evolutionary Genetics is funded by the French “Investissement d’avenir” program, Laboratoires d’Excellence IBEID (grant # ANR-10-LABX-62-IBEID), and the FRM (Equipe FRM DEQ20180339214).

## Methods

### Exome sequencing data

Human genetic variants from the Exome Aggregation Consortium (ExAC) database (12) (http://exac.broadinstitute.org/) were downloaded on September 9^th^ 2016, release 0.3.1, non-TCGA subset. Variants from Exome data of the Genome Aggregation Database (gnomAD, http://gnomad.broadinstitute.org/) were obtained on February 28^th^ 2017, release 2.0.1 ExAC, gnomAD variants were annotated with the Ensembl Variant Effect Predictor VEP (70) (v81 for ExAC, v85 for gnomAD), with loss-of-function (LoF) annotations from LOFTEE (https://github.com/konradjk/loftee). Homo sapiens genome build GRCh37/hg19 was used with both databases. Analyses were restricted throughout this study to a background set of 20232 human protein-coding genes obtained from BioMart Ensembl 75, version Feb 2014 (GRCh37.p13; (71)).

Protein-coding genes are defined as those containing an open reading frame (ORF). On the contrary, pseudogenes are typically defined as gene losses resulting from fixations of null alleles that occurred in the human lineage after a speciation event, some of them may actually be human-specific, i.e. fixed after the human-chimpanzee divergence. However, the definition might be larger, including the so-called processed pseudogenes, corresponding to DNA sequences reverse-transcribed from RNA and randomly inserted into the genome (62, 63). A total of 20,232 protein-coding genes and 13,921 pseudogenes are reported by Ensembl following the Ensembl Genebuild workflow incorporating the HAVANA group manual annotations (27). Among the protein-coding genes, 382 genes were identified with a gene name starting with “Olfactory receptor”.

### Loss-of-function variants

Loss-of-function (LoF) variants are considered here as being those predicted to lead to an early stop-gain, indel frameshift or essential splice-site disruption (i.e, splice-site donor and splice-site acceptor variants). The following low-confidence LoF variants were filtered out: variants where the purported LoF allele is the ancestral state (across primates), stop-gain and frameshift variants in the last 5% of the transcript, or in an exon with non-canonical splice sites around it (i.e. intron does not start with GT and end with AG), and splice-site variants in small introns (<15 bp), in an intron with a non-canonical splice site or rescued by nearby in-frame splice sites. Following the criteria used in the ExAC flagship paper (12), only LOFTEE-labeled “high confidence” LoF variants mapped to canonical isoforms, with a genotype quality ≥ 20, depth ≥ 10, and a call rate > 80% were kept. Following https://macarthurlab.org/2016/03/17/reproduce-all-the-figures-a-users-guide-to-exac-part-2/, variants in the top 10 most-multiallelic kilobases of the human genome were filtered out, i.e: Chr14:106330000-106331000; Chr2:89160000-89161000; Chr14:106329000-106330000; Chr14:107178000-107179000; Chr17:18967000-18968000; Chr22:23223000-23224000; Chr1:152975000-152976000; Chr2:89161000-89162000; Chr14:107179000-107180000; Chr17:19091000-19092000. For GnomAD variants, a heterozygote genotype allele balance > 0.2 was further required.

### Definition of the set of dispensable genes

In addition to the previous criteria, a set of filters for LoF variants was adopted in this work retaining variants with a Variant Quality Score Recalibration (VQSR) equal to ‘PASS’ both for single nucleotide polymorphisms (SNPs) and frameshifts and affecting the APPRIS principal isoform (28) (downloaded on downloaded March 2nd 2017, using Gencode19/Ensembl74). Dispensable genes were defined as those presenting an aggregated frequency of homozygous individuals (all LoF variants combined) higher than 1% in at least one of the 5 main populations considered, i.e.: Africans (including African Americans), East Asians, South Asians, Europeans (Finnish and Non-Finnish), and Americans.

### Impact prediction of LoF variants

The mRNA region capable of triggering transcript degradation by NMD upon an early stop-gain was defined as previously (5) following HAVANA annotation guidelines (v.20). Specifically, the NMD-target region of a transcript was defined as those positions more than 50 nucleotides upstream the 3’-most exon-exon junction. Transcripts bearing stop-gain variants at these regions are predicted to be degraded by NMD (72). Molecular impact prediction of splice-disrupting variants was performed using the Human Splicing Finder (HSF) software ((73); online version 3.1 available at http://www.umd.be/HSF/ with default parameters). HSF classifies splicing variants into 5 categories: unknown impact, no impact, probably no impact, potential alternation, most probably affecting variant. LoF variants were classified into those predicted to be LoF with low and higher probability. Low probability LoF arbitrarily include (i) stop-gains and frameshift variants truncating the last 15% of the protein sequence, which might translate in small truncations in the final protein, or mapping into the first 100 nucleotides of the transcript, which has been reported as a window length where LoF variants are often recovered by alternative start sites (74); and (ii) essential splice site variants with an unknown or low computationally predicted impact on splicing motifs based on position weight matrices, maximum entropy and motif comparison methods (Methods). High probability LoF variants include (i) stop-gains and frameshifts truncating more than 15% of the protein sequence or occurring in a region prone to transcript degradation by NMD, which probably result in complete abrogation of protein production (Methods); and (ii) putative splice-site variants with an intermediate or high computationally predicted impact (Methods). All associations of LoF variants reported in the GWAS catalog (version v1.0, date 2019-01-11) were downloaded from ftp://ftp.ebi.ac.uk/pub/databases/gwas/.

### Gene Ontology enrichment analysis

Gene Ontology enrichment analysis was performed with topGO R package from Bioconductor (version 3.5), using Fisher’s Exact test option with parameters p-value threshold < 0.005 and number of significant genes found with GO term ≥ 5.

### Gene expression patterns across human organs and immune cell types

Two lists of organ and tissue-expressed genes were defined based on the RNA-seq expression data from the Illumina Body Map project (IBM, 15,688 expressed genes), and the GTEX project (16,762 expressed genes). First, RNA-seq expression data from a panel of 11 human organs and tissues (one sample each) from the Illumina Body Map project (IBM) was taken from the Expression Atlas database (EBI accession E-MTAB-513; 1 February 2017 release; https://www.ebi.ac.uk/gxa/experiments/E-MTAB-513/Results). The list of organs and tissues included: adipose tissue, brain, breast, colon, heart, kidney, liver, lung, ovary, skeletal muscle tissue, and testis. It should be noted that leukocyte, lymph node, adrenal, prostate and thyroid gland data were removed from these datasets. A total of 15,688 organ-expressed genes were defined as being expressed in more than 3 Transcripts Per Million (TPM; following (75)) in at least one of the IBM samples considered. Second, RNA-seq expression data from a panel of 24 human organs (multiple samples per organ) from the GTEX project (76) was taken from https://gtexportal.org/home/tissueSummaryPage (version V6p). Blood, blood vessel, salivary gland, adrenal gland, thyroid, pituitary and bone marrow were removed from these datasets. A total of 16,762 organ-expressed genes were defined as being expressed in more than 3 TPM in at least 20% of the samples from at least one GTEX organ. The organ-expressed set of genes previously defined was further classified into a set of “organ-specific genes” and “organ-pervasive genes”, depending on whether the gene was defined as expressed in <20% (organ-specific) or >80% of the organs and tissue types evaluated in the corresponding dataset (i.e. IBM or GTEX).

In addition, the RNA-seq data generated by the BluePrint project (77) was used as a reference set of gene expression for the different immune cell types from human venous blood origin (August 19, 2017 release, data available at http://dcc.blueprint-epigenome.eu/#/files). Only cell types with more than 2 samples were considered. A total of 85 libraries were retained including: 9 B cell and 17 T cell samples (collectively considered as adaptive immune cell types), and 15 monocytes, 25 macrophages, 6 dendritic cell and 13 neutrophil samples (collectively considered as innate immune cell types). A total of 7,346 adaptive immune cell-expressed genes were detected as those expressed in more than 3 TPM in B cell or T cells in at least in 20% of the corresponding samples collectively considered. Analogously, a total of 9,069 innate immune cell-expressed genes were defined here as those expressed in more than 3 TPM in macrophages, monocytes, neutrophils, or dendritic cells in at least 20% of the corresponding samples collectively considered. Full details about the libraries used as provided by the BluePrint project are reported in **Supplementary Table 7**. The set of genes not found to be expressed in any of the previous lists was determined from the complement of the reference list of 20120 protein-coding genes defined as Ensembl Biomart, release 75 (71).

### Gene-level annotations

The following gene-level features associated with natural selection were obtained: Gene Damage Index (GDI) scores, a gene-level metric of the mutational damage that has accumulated in the general population, based on CADD scores, were taken from (44). High GDI values reflect highly damaged genes. The Residual Variation Intolerance Score (RVIS percentile, provided in (45)), assesses the gene departure from the average number of common functional mutations in genes with a similar amount of mutational burden in humans. High RVIS percentiles reflect genes that are highly tolerant to variation. The gene probability of loss-of-function intolerance (pLI, (12)), estimating the depletion of rare and *de novo* protein-truncating variants compared to the expectations derived from a neutral model of de novo variation on ExAC exomes data, as well as pRec, estimating gene intolerance to two rare and *de novo* protein-truncating variants (analogous to recessive genes) were obtained from the ExAC Browser (release 0.3.1, (46)). pLI and pRec values close to 1 represent gene intolerance to heterozygous and homozygous loss-of-function and to homozygous mutations, respectively. “f” values were obtained through SnIPRE (48) following (47), based on the comparison of polymorphism and divergence at synonymous and non-synonymous sites. Data on the number of human paralogs for each gene were collected from the OGEE database (78). Human essential genes were obtained from (33). Following the original authors, the top 15% percentile of *in-vivo* and *in-vitro* human essential genes and the complete list of knockout mouse genes were retained.

Monogenic Mendelian disease genes were obtained following Chong et al. (79): OMIM raw data files were downloaded from (80). Phenotype descriptions containing the word ‘somatic’ were flagged as ‘somatic’, and those containing ‘risk’, ‘quantitative trait locus’, ‘QTL’, ‘{’, ‘[’ or ‘susceptibility to’ were flagged as ‘complex’. Monogenic Mendelian genes were defined as those having a supporting evidence level of 3 (i.e. the molecular basis of the disease is known) and not having a ‘somatic’ or ‘complex’ flag.

### Genome-wide scans for recent positive selection at loss-of-function mutations

To test for the occurrence of positive selection of loss-of-function mutations we computed two neutrality statistics: the interpopulation *F*_ST_ that identifies loci exhibiting high variation in allele frequencies between groups of populations (49), and the intrapopulation integrated haplotype score (iHS) (50), that compares the extent of haplotype homozygosity at the ancestral and derived alleles. Positive selection analyses were confined to biallelic SNPs found in the 1000 Genomes Project phase 3 data (81), including 2,504 individuals from 26 populations, assigned to 5 meta-populations and predicted to have severe damaging consequences. (**Supplementary Table 8**). Multiallelic SNPs, SNPs not detected in the 1000 Genomes Project and indel frameshifts were discarded in the positive selection analysis. For *F*_ST_ calculation, we investigated a total of 78 LoF mutations that passed quality filters, and compared the allele frequencies of these variants in 26 populations, to the allele frequencies of the same mutations in the European (CEU) and African (YRI) reference populations (**Supplementary Table 8**). More specifically, we compared allele frequencies in populations from the African (AFR), East Asian (EAS) and South Asian (SAS) meta-populations to allele frequencies in the CEU groups, and allele frequencies in populations from the American (AMR) and European (EUR) meta-populations to allele frequencies in the YRI group. To detect candidate variants for positive selection based on *F*_ST_ values, we used an outlier approach and considered LoF mutations presenting *F*_ST_ values located in the top 5% of the distribution of *F*_ST_ genome-wide. We identified 32 LoF mutations presenting high *F*_ST_ values in at least one population in 32 genes (including 8 mutations located in OR genes). For haplotype-based iHS score calculations, we first defined the derived allele state of each SNP based on the 6-EPO alignment, and retained only SNPs with a derived allele frequency (DAF) between 10% and 90% to maximize the power of iHS to detect selective signals. These additional filters led to a total of 72 LoF mutations to be investigated. We calculated iHS scores in 100kb windows with home-made scripts and normalized values. To detect selection events targeting derived alleles, we considered LoF mutations located in the top 5% of most negative iHS values genome-wide and found a total of 15 LoF mutations with iHS scores in the top 5% of lowest iHS values genome wide in at least one population (including 1 mutation located in an OR gene).

## Supplementary Figures

**Supplementary Figure 1.**
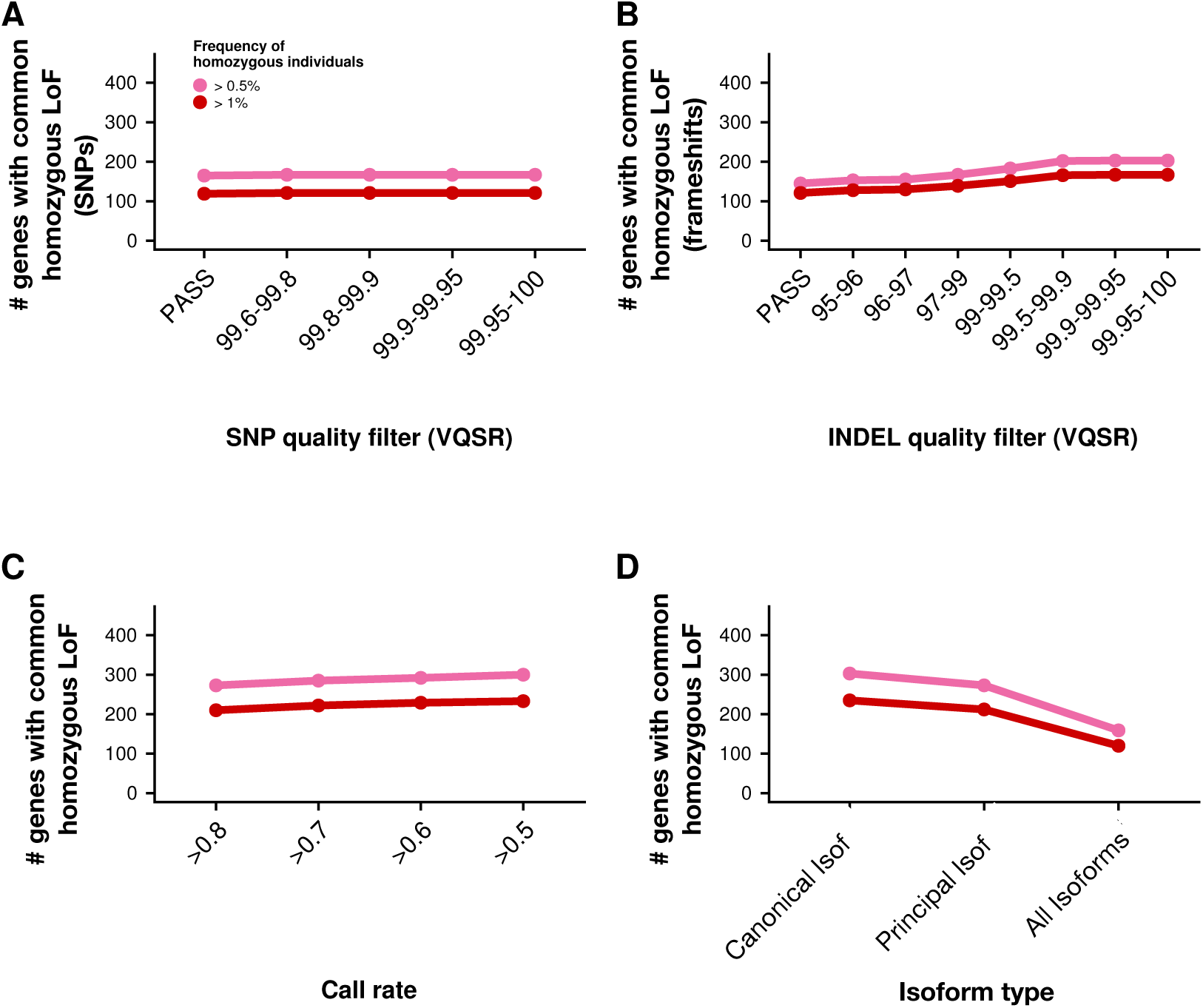
Impact of the different filtering criteria on the final number of dispensable genes detected in the ExAC database. **A.** Number of genes with common homozygous LoF caused by Single Nucleotide Polymorphisms as a function of the variant quality score recalibration (VQSR) threshold. **B.** Number of genes with homozygous LoF caused by frameshifts as a function of the variant quality score recalibration (VQSR) threshold. **C.** Number of genes with common homozygous LoF caused by SNPs and frameshifts as a function of the call rate threshold. **D**. Number of genes with common homozygous LoF caused by SNPs and frameshifts depending on whether LoF variant affects any canonical isoform, only the principal isoform or all canonical isoforms (variant constitutive of all isoforms; **Methods**).

**Supplementary Figure 2.**
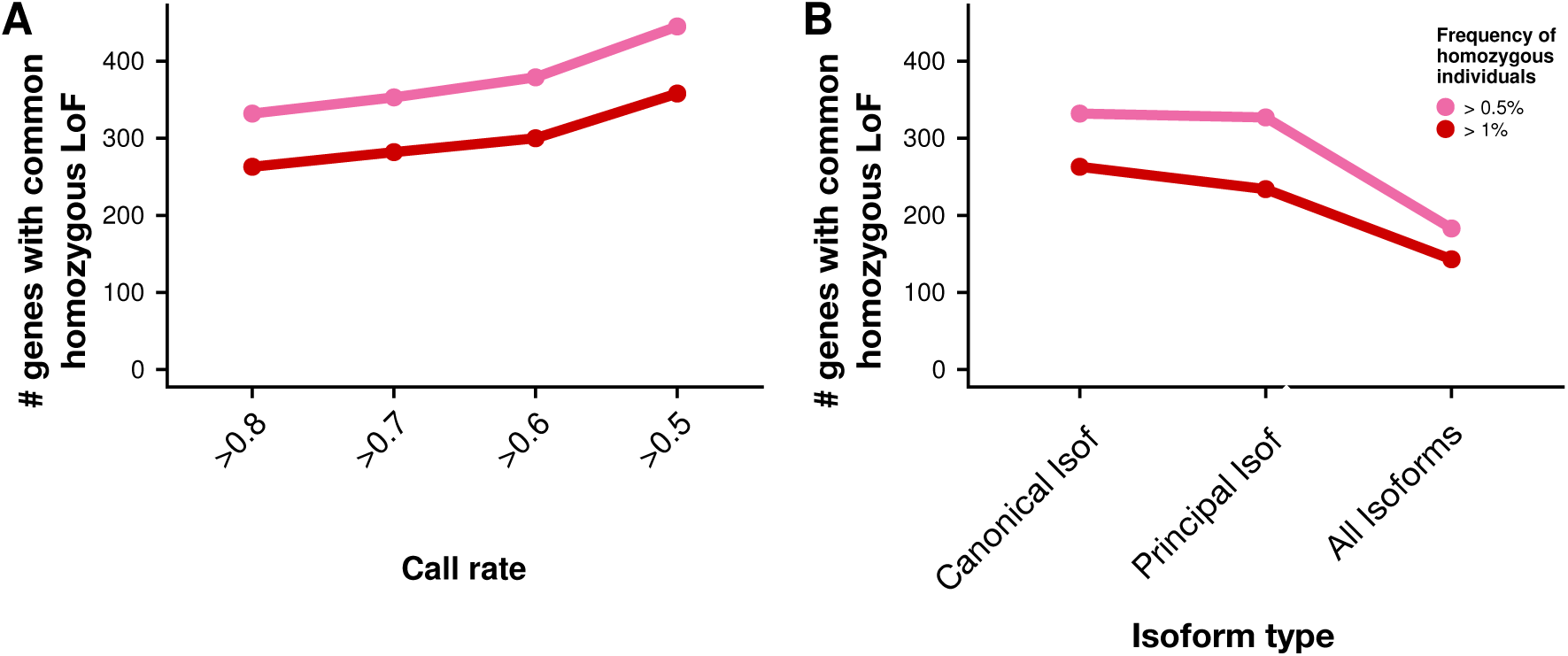
Impact of the different filtering criteria adopted on the final number of dispensable genes detected in the GnomAD database. **A**. Number of genes with common homozygous LoF caused by SNPs and frameshifts as a function of the call rate threshold. **B**. Number of genes with common homozygous LoF caused by SNPs and frameshifts depending on whether the LoF variant affects the canonical isoform, the principal isoform or all isoforms (variant constitutive of all isoforms). It should be noted that VQSR scores were not used in the GnomAD database, thus panels analogous to **Supplementary Figure 1 A** and **B** could not be drawn.

**Supplementary Figure 3.**
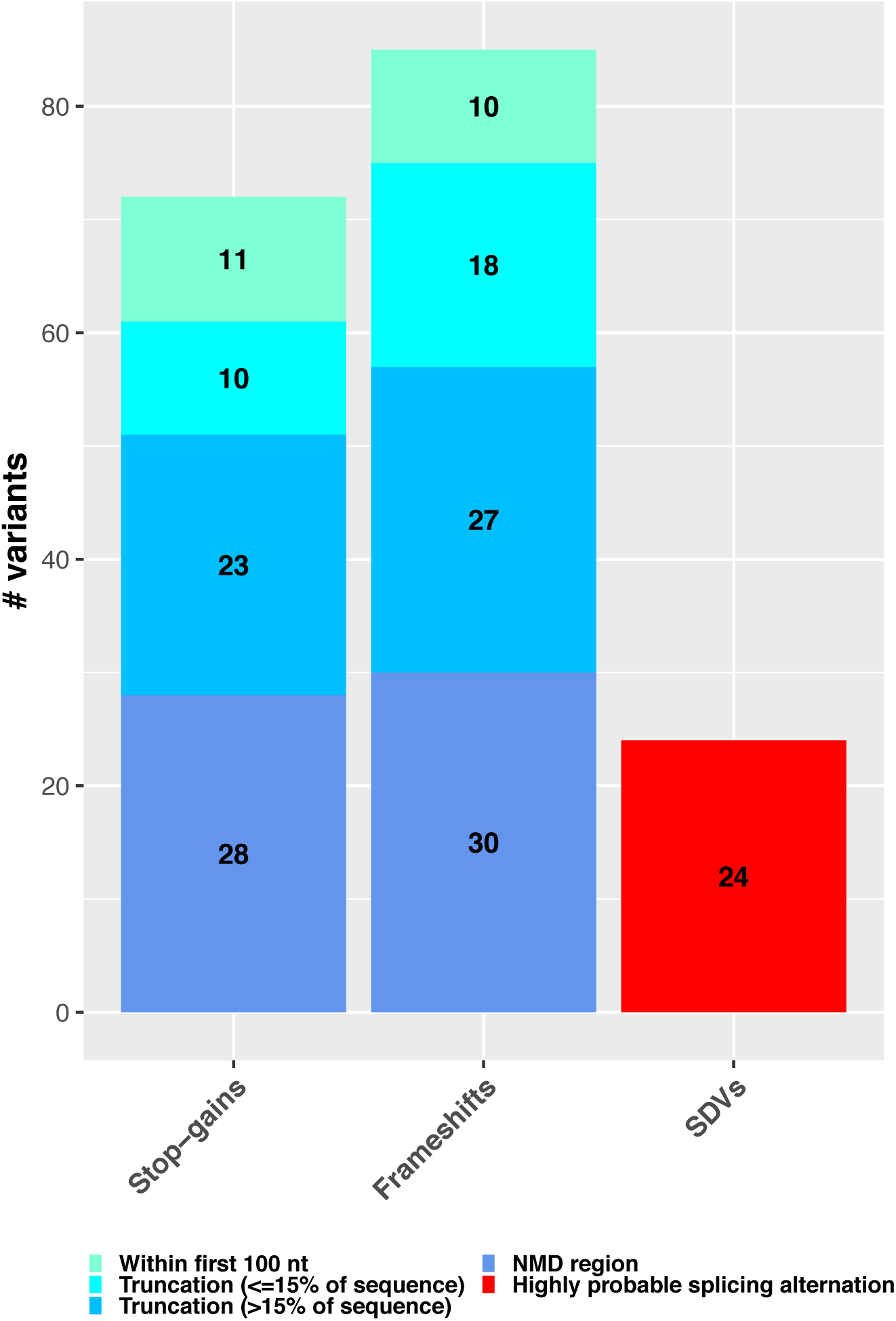
Predicted functional impact of LoF variants defining the set of dispensable protein coding genes. Bar plots show the distribution of LoF variants that define the set of dispensable genes according to their molecular consequences (stop-gains, frameshifts and splice-disrupting variants, SDV) and the predicted type of functional impact, according to the following categories: In the case of stop-gains and frameshifts, LoF variants are classified among those i) mapping to the first 100 nucleotides of the associated transcript, ii) potentially triggering NMD, or iii) truncating more, or less or equal than 15% of the affected protein sequence. In the case of putative splice-disrupting variants (SDVs), severity was computationally predicted as unknown, very low, intermediate or high impact (**Methods**).

**Supplementary Figure 4.**
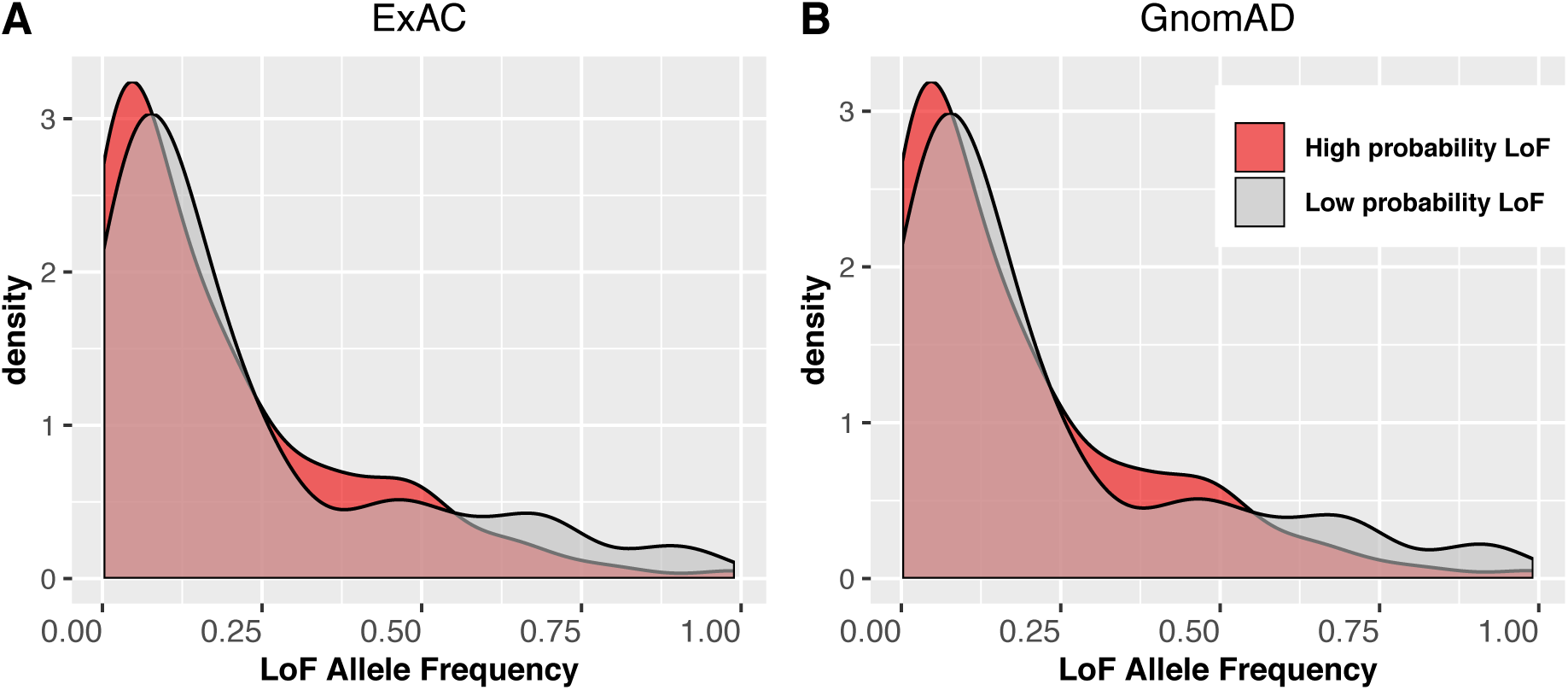
Allele Frequency distribution of LoF variants defining the set of dispensable protein-coding genes. Allele Frequency of LoF variants is represented separately for low probability LoF (light grey) and high probably LoF (dark red) variants. Allele frequencies from ExAC were used in **Panel A**, whereas GnomAD data were used in **Panel B**.

**Supplementary Figure 5.**
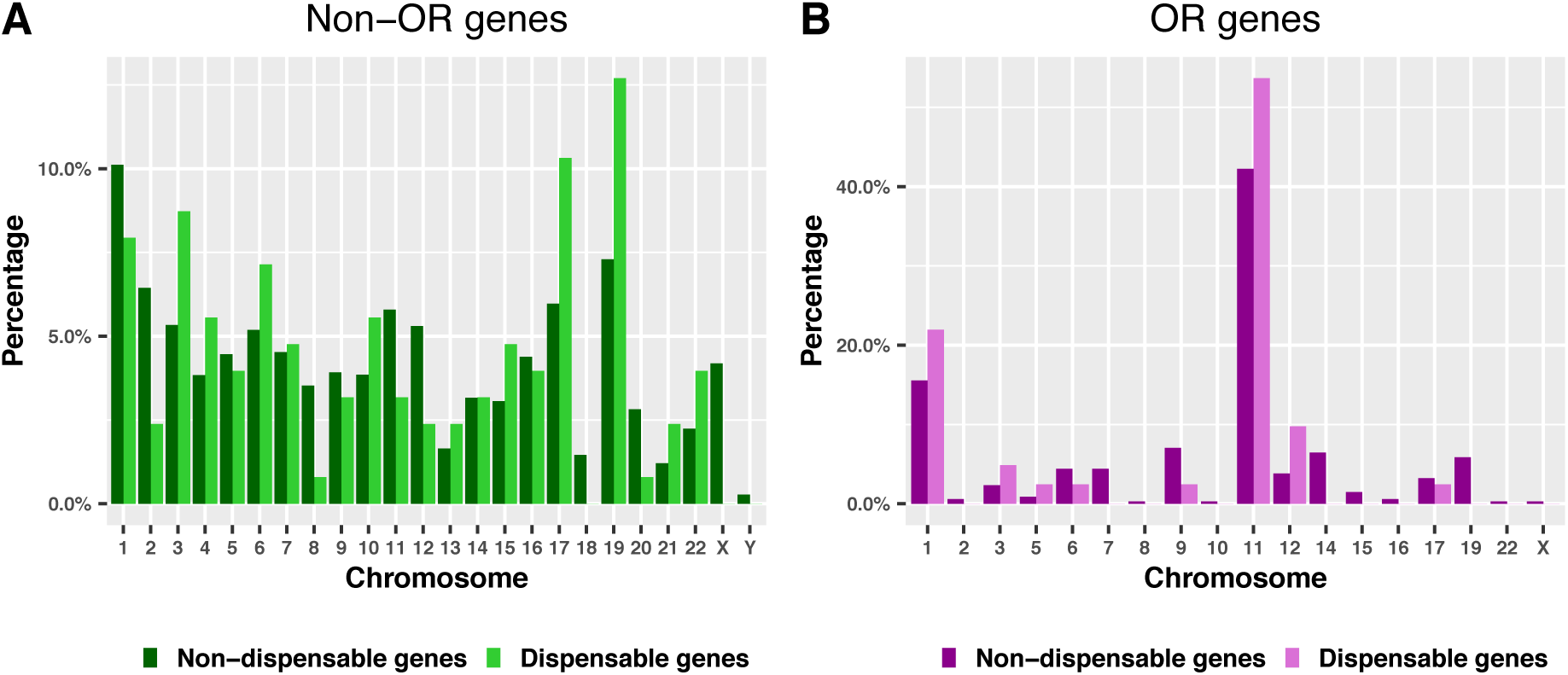
Distribution of dispensable and non-dispensable genes across chromosomes. Barplots display the percentage of genes across human chromosomes of the following 4 gene sets, each adding to 100%: (**A**) dispensable non-OR genes (light green) and non-dispensable non-OR genes (dark green), and (**B**) dispensable OR genes (light purple) and non-dispensable OR genes (dark purple).

**Supplementary Figure 6.**
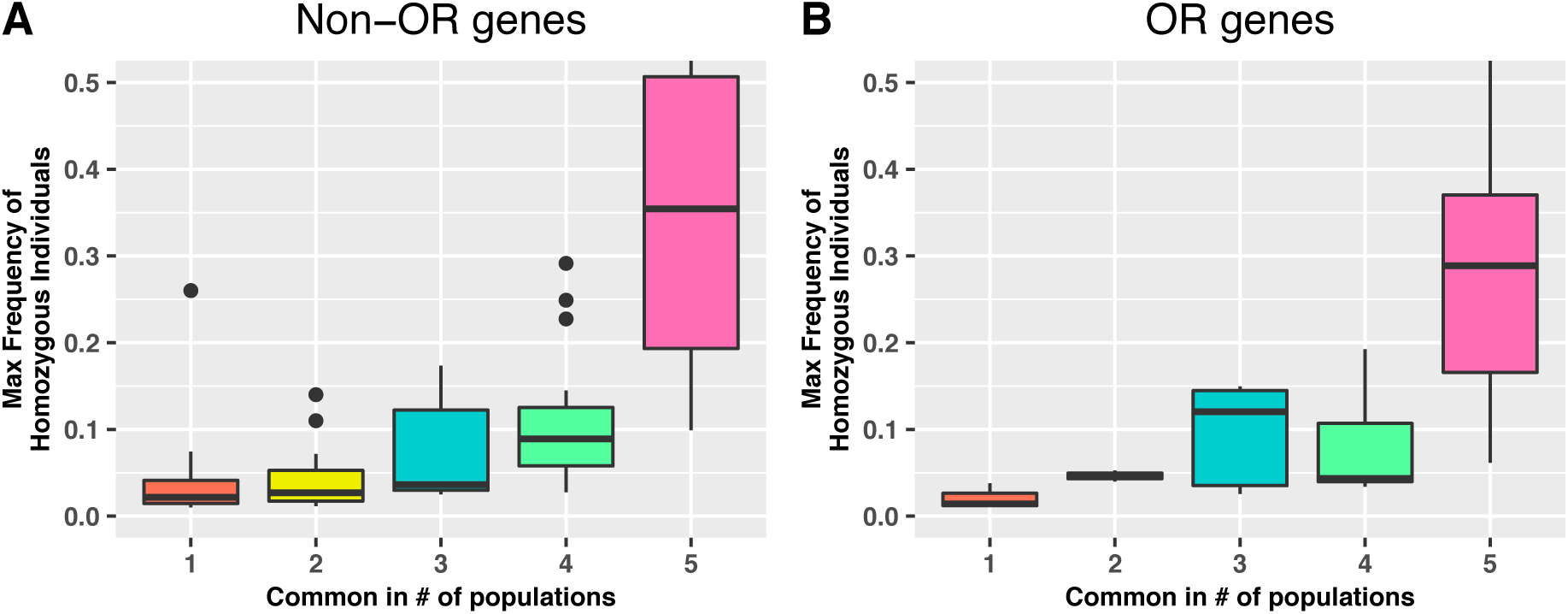
Distribution of the maximum frequency of homozygous individuals for dispensable non-OR genes (**A**) and dispensable OR genes (**B**) as a function of the number of populations in which they were found to be dispensable. As in Figure 3, the homozygous LoF variant frequencies were taken from the GnomAD dataset.

**Supplementary Figure 7.**
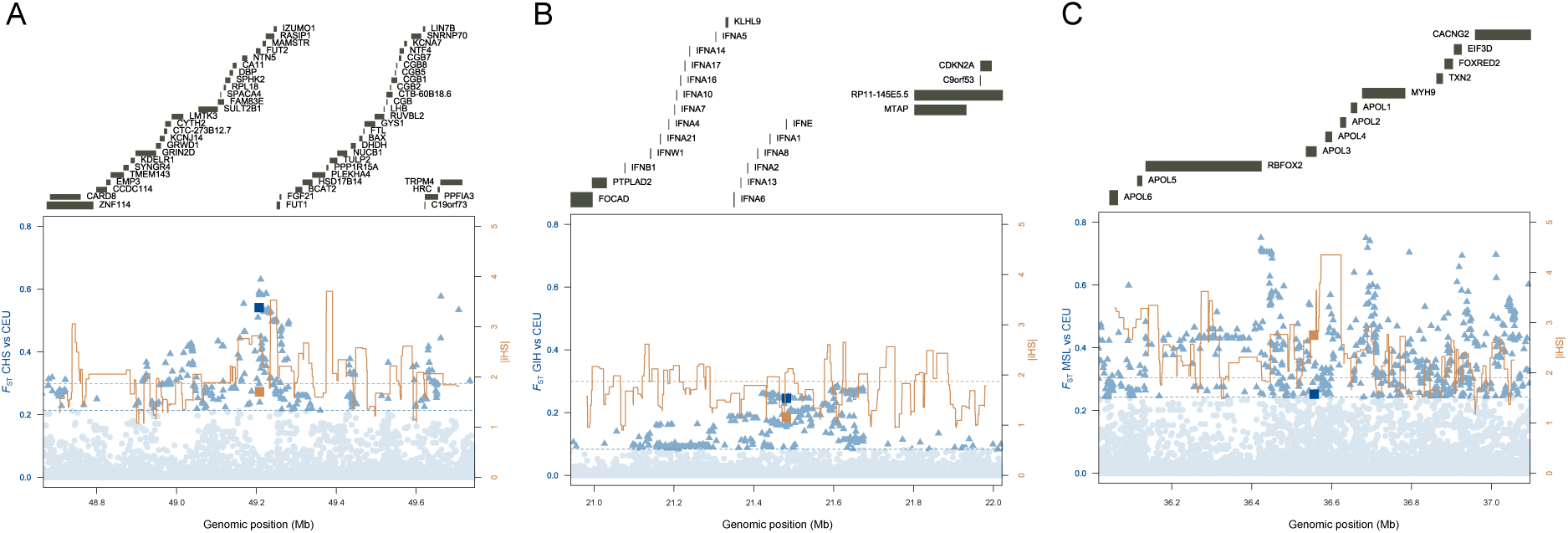
Selective sweep signals. Local genomic signatures of positive selection in 1Mb regions around LoF mutations located in (**A**) *FUT2* (chr19:49206674) for CEU, (**B**) *IFNE* (chr9:21481483) for GIH and (**C**) *APOL3* (chr22:36556768) for MSL. Blue and orange squares indicate *F*_ST_ and |iHS| values respectively at the LoF allele. Blue dots and triangles indicate SNP *F*_ST_ percentiles and the blue dashed line indicate 95^th^ percentile of *F*_ST_ values genome-wide. Orange solid line indicate the maximum |iHS| value in sliding windows of 50 SNPs and the orange dashed line indicate the 95^th^ percentile of |iHS| values genome-wide.

**Supplementary Figure 8.**
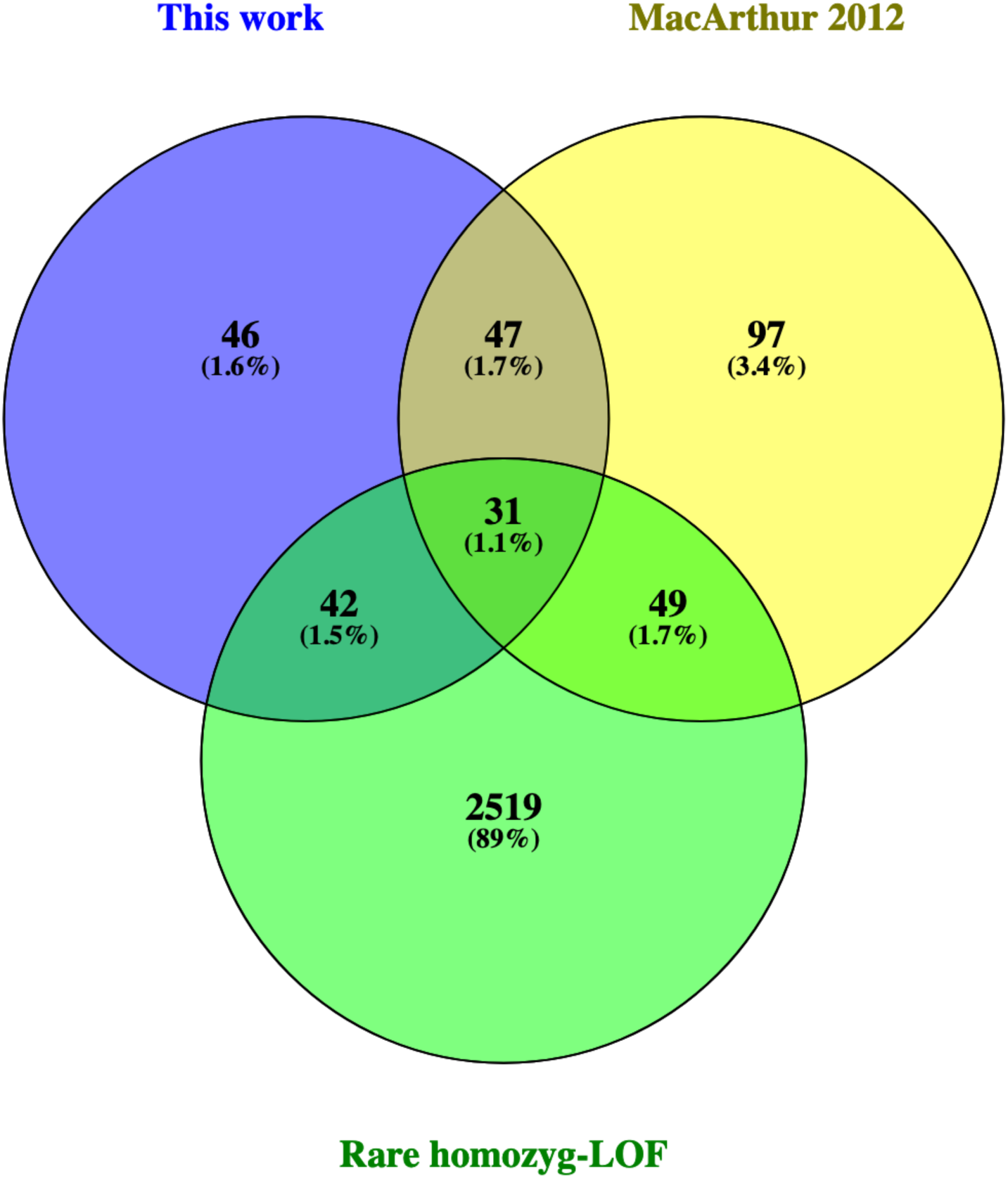
Overlap of the dispensable genes detected in this work with those identified in previous studies. The figures show the Venn diagrams representing the overlap of the 166 putatively dispensable genes detected in this work with 253 genes apparently tolerant to homozygous rare LoF variants reported in MacArthur et al. (9) and a total list of 2641 presenting homozygous rare LoF variants reported from bottlenecked or consanguineous populations (10,11,13,14).

## Supplementary Tables

**Supplementary Tables 1-7 in an attached excel file**

**Supplementary Table 8.** Populations from the 1000 Genomes Project used in the positive selection analysis of common LoF variants.

| Population Code | Population Description | Sample_size | Super Population Code |
| --- | --- | --- | --- |
| ACB | African Caribbeans in Barbados | 96 | AFR |
| ASW | Americans of African Ancestry in SW USA | 61 | AFR |
| ESN | Esan in Nigeria | 99 | AFR |
| GWD | Gambian in Western Divisions in the Gambia | 113 | AFR |
| LWK | Luhya in Webuye, Kenya | 99 | AFR |
| MSL | Mende in Sierra Leone | 85 | AFR |
| YRI | Yoruba in Ibadan, Nigeria | 108 | AFR |
| CLM | Colombians from Medellin, Colombia | 94 | AMR |
| MXL | Mexican Ancestry from Los Angeles USA | 64 | AMR |
| PEL | Peruvians from Lima, Peru | 85 | AMR |
| PUR | Puerto Ricans from Puerto Rico | 104 | AMR |
| CDX | Chinese Dai in Xishuangbanna, China | 93 | EAS |
| CHB | Han Chinese in Beijing, China | 103 | EAS |
| CHS | Southern Han Chinese | 105 | EAS |
| JPT | Japanese in Tokyo, Japan | 104 | EAS |
| KHV | Kinh in Ho Chi Minh City, Vietnam | 99 | EAS |
| CEU | Utah Residents (CEPH) with Northern and Western European Ancestry | 99 | EUR |
| FIN | Finnish in Finland | 99 | EUR |
| GBR | British in England and Scotland | 91 | EUR |
| IBS | Iberian Population in Spain | 107 | EUR |
| TSI | Toscani in Italia | 107 | EUR |
| BEB | Bengali from Bangladesh | 86 | SAS |
| GIH | Gujarati Indian from Houston, Texas | 103 | SAS |
| ITU | Indian Telugu from the UK | 102 | SAS |
| PJL | Punjabi from Lahore, Pakistan | 96 | SAS |
| STU | Sri Lankan Tamil from the UK | 102 | SAS |

